# Tracking SARS-CoV-2 Spike Protein Mutations in the United States (2020/01 – 2021/03) Using a Statistical Learning Strategy

**DOI:** 10.1101/2021.06.15.448495

**Authors:** Lue Ping Zhao, Terry P. Lybrand, Peter B. Gilbert, Thomas R. Hawn, Joshua T. Schiffer, Leonidas Stamatatos, Thomas H. Payne, Lindsay N. Carpp, Daniel E. Geraghty, Keith R. Jerome

**Affiliations:** Public Health Sciences Division, Fred Hutchinson Cancer Research Center; Seattle, WA, USA; Quintepa Computing LLC; Nashville, TN, USA; Department of Chemistry; Department of Pharmacology, Vanderbilt University; Nashville, TN, USA; Vaccine and Infectious Disease Division, Fred Hutchinson Cancer Research Center; Seattle, WA, USA; Department of Medicine, University of Washington School of Medicine; Seattle, WA, USA; Department of Global Health, University of Washington; Seattle, WA, USA; Clinical Research Division, Fred Hutchinson Cancer Research Center, Seattle; WA, USA

## Abstract

The emergence and establishment of SARS-CoV-2 variants of interest (VOI) and variants of concern (VOC) highlight the importance of genomic surveillance. We propose a statistical learning strategy (SLS) for identifying and spatiotemporally tracking potentially relevant Spike protein mutations. We analyzed 167,893 Spike protein sequences from US COVID-19 cases (excluding 21,391 sequences from VOI/VOC strains) deposited at GISAID from January 19, 2020 to March 15, 2021. Alignment against the reference Spike protein sequence led to the identification of viral residue variants (VRVs), i.e., residues harboring a substitution compared to the reference strain. Next, generalized additive models were applied to model VRV temporal dynamics, to identify VRVs with significant and substantial dynamics (false discovery rate q-value <0.01; maximum VRV proportion > 10% on at least one day).

Unsupervised learning was then applied to hierarchically organize VRVs by spatiotemporal patterns and identify VRV-haplotypes. Finally, homology modelling was performed to gain insight into potential impact of VRVs on Spike protein structure. We identified 90 VRVs, 71 of which have not previously been observed in a VOI/VOC, and 35 of which have emerged recently and are durably present. Our analysis identifies 17 VRVs ∼91 days earlier than their first corresponding VOI/VOC publication. Unsupervised learning revealed eight VRV-haplotypes of 4 VRVs or more, suggesting two emerging strains (B1.1.222 and B.1.234). Structural modeling supported potential functional impact of the D1118H and L452R mutations. The SLS approach equally monitors all Spike residues over time, independently of existing phylogenic classifications, and is complementary to existing genomic surveillance methods.

## INTRODUCTION

Severe Acute Respiratory Syndrome coronavirus 2 (SARS-CoV-2), the pathogen responsible for the global Covid-19 pandemic, is an RNA virus and thus prone to replication errors (*1*). Replication errors that yield nonsynonymous amino acid (AA) substitutions, or nucleotide insertions or deletions that cause a frame shift and alter the subsequent coding sequence, can lead to a variety of outcomes. If the resulting mutations have detrimental effects on fitness, or if they have neutral effects on fitness and undergo stochastic extinction, variants harboring these mutations fail to become established in the population. However, mutations that confer a fitness advantage can rapidly become dominant in a population. For SARS-CoV-2, there are three classes of variant: Variant of Interest (VOI), Variant of Concern (VOC), and Variant of High Consequence (VOHC). The CDC is currently monitoring and characterizing 8 VOIs (B.1.526, B.1.526.1, B.1.525, P.2, B.1.617, B.1.617.1, B.1.617.2, B.1.617.3) and 5 VOCs (B.1.1.7, P.1, B.1.351, B.1.427, B.1.429) in the United States (*2*). VOCs show specific attributes such as increased transmissibility (*3-7*), increased resistance to neutralization by antibodies elicited through natural infection (*3, 8-10*), and/or increased resistance to neutralization by vaccine-elicited antibodies (*10-12*), and have already influenced vaccine development, evidenced by the current planning of clinical trials to test variant-adapted vaccines (*13*). While no VOHCs have yet been identified, it remains possible that such variants – i.e. variants that can effectively evade natural or vaccine-induced immunity – may yet emerge (*14, 15*). The identification of VOHCs could necessitate the introduction of more stringent public health guidelines and/or spur further treatment and vaccine development.

Genomic surveillance is critical for tracking the emergence and spread of new variants. Such surveillance can be accomplished via a variety of approaches, such as phylogenic analysis (*3, 16*). In this approach, new viral sequences are classified to existing lineages identified by PANGO (*17*), subsets of samples with the same branches are identified, and variant frequencies are counted to identify new variants. The NextStrain methodology (*18*) can model dynamic changes of variant proportions, while an alternative approach aligns sequence data to a matrix of binary indicators for the presence of variants, and systematically evaluates each mutant as a potential variant (*19*). Leveraging the analytic approach of single nucleotide polymorphisms (SNPs), variants have been identified by assessing linkage-disequilibrium (*20*) or similar SNP-based identification and analysis (*21*). However, with the exception of the NextStrain methodology (*3*), these methods do not directly take into account sequence collection time, nor explicitly incorporate highly granular geographic information. Moreover, these methods take a holistic view of the viral genome. Thus, there is a need for complementary approaches for detecting and characterizing Spike mutations of potential public health importance that may be missed, or detected later, by existing genomic surveillance methods.

To meet this need, we describe a statistical learning strategy (SLS) using generalized additive models, unsupervised learning techniques, and single nucleotide polymorphism (SNP) methodologies for identifying and spatiotemporally characterizing viral residue variants (VRVs), a term we use to describe AA positions in the Spike protein where a mutation is significantly present in a given geographic area. The SLS method generates pertinent statistics for reproducible scientific inference and facilitates visual representation of results for intuitive interpretation. Using publicly available SARS-CoV-2 sequences from US COVID-19 cases that were not assigned to a VOI or VOC lineage, we apply our method to identify and spatiotemporally characterize, within individual US states/territories, VRVs in the Spike protein. We also apply standard homology modeling methods to highlight individual AA mutations with the potential to impact Spike protein structure and/or function.

## RESULTS

### Ab Initio Discovery of VRVs

We first applied the SLS method to identify VRVs separately in each state/territory (Fig. S1). The decision to compartmentalize VRV discovery by state/territory was partially based on the fact that domestic travel restrictions have varied over the course of the pandemic, with nearly half of all states having imposed some type of interstate travel restriction (*22*), leading to the hypothesis that VRVs may follow state/territory-specific temporal dynamics. The identified VRVs showed a range of dynamic patterns across the different states/territories (Fig. S2), exemplified by the five different trajectories taken by the V382, L452, T478, P681, and T732 VRVs in California (Fig. 1A). The relative abundance of V382 started rising on day 250, exceeded 10% on day 259, and fell below 10% on day 275. L452 emerged on day 310, exceeded 10% on day 390, and exhibited a positive trajectory thereafter. Three other VRVs (T478, P681, T732) had similar trajectories to L452.

**Fig. 1.**
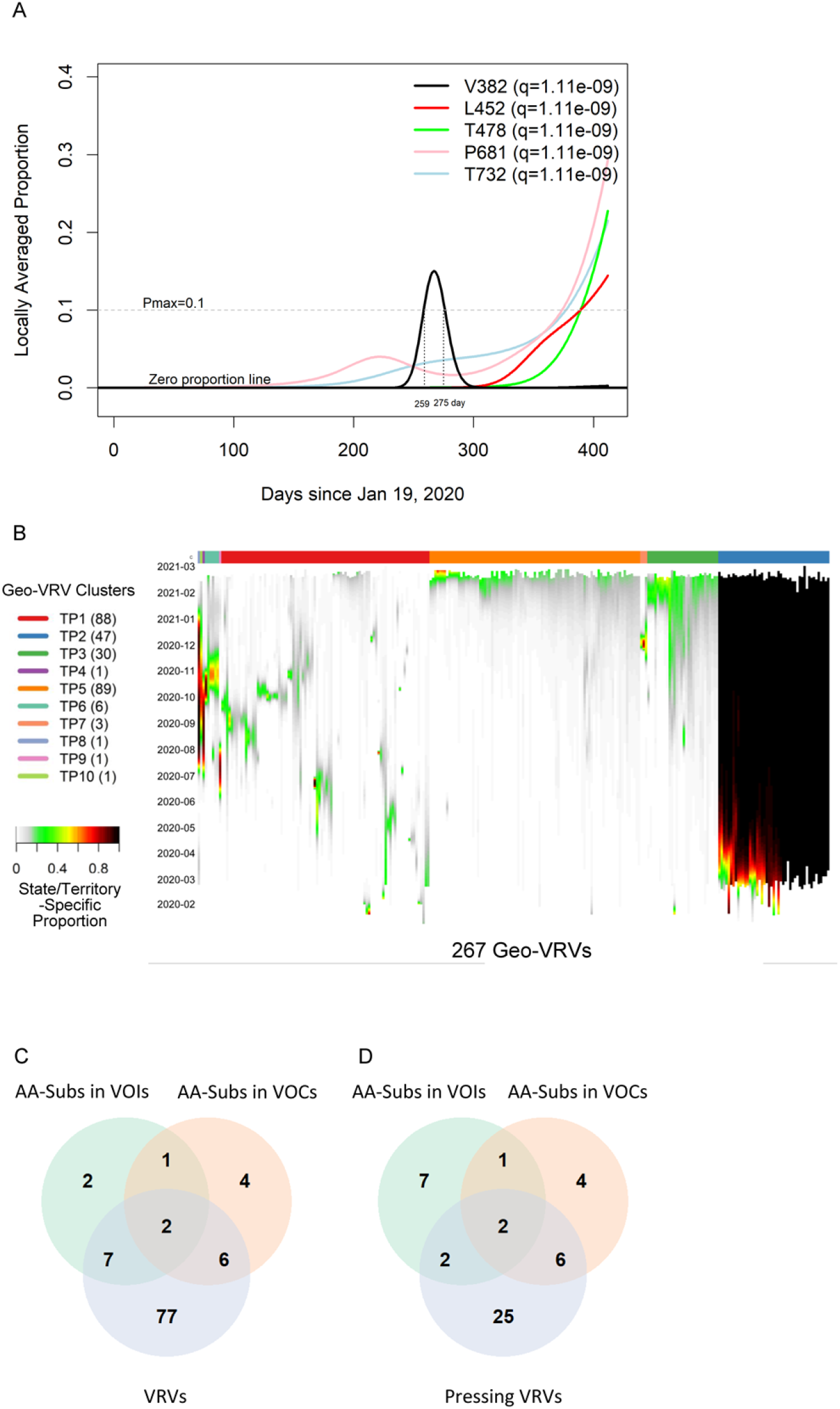
Viral Residue Variant (VRV) spatiotemporal patterns in the United States. (**A**) Locally averaged proportions over time of five VRVs (V382, L452, T478, P681 and T732), modeled using sequences from California. The horizontal gray dotted line denotes the Pmax cutoff of 10%. V382 exceeded the Pmax cutoff of 10% on day 259, and dropped below the Pmax cutoff of 10% on day 275 (marked by the vertical gray lines). (**B**) Heatmap of the 267 identified geo-VRVs, with color designating the state/territory-specific VRV proportion at the sampling time as designated on the left-hand vertical axis. Geo-VRVs with similar temporal dynamics are grouped into 10 clusters (TP1 through TP10), as designated by the color bar at the top of the heatmap. (**C, D**) Venn diagrams showing the relationships between AA-subs in VOIs, AA-subs in VOCs, and (**C**) VRVs or (**D**) pressing VRVs. AA-subs, amino acid positions that have been shown to harbor substitutions within US-circulating variants; VOCs, variants of concern; VOIs, variants of interest.

We refer to the combination of a VRV and a state/territory in which it was identified as a “geo-VRV”. A total of 267 geo-VRVs, consisting of combinations of 90 VRVs identified among the 52 state/territory classifications, were identified (Table S3). Fifty-eight VRVs were only observed in one state/territory, whereas 32 were observed in two or more (Table S4).

Unsupervised learning was next applied to organize the 267 geo-VRVs into 10 clusters (TP1 through TP10) (Fig. 1B, Table S3). The cluster most strikingly different from the others was “TP2”, which was composed of 47 geo-VRVs, each of which contained the D614 VRV at a maximum relative abundance of 100%, showing the early dominance of the D614 VRV in these states/territories. Clusters TP3, and TP5 include geo-VRVs of potential concern, since they include VRVs that appear to have emerged within the last few months in their specific states/territories. In contrast, most VRVs in the remaining clusters tended to expand and contract within relatively short times in a given state/territory, making such VRVs likely less important from a public health perspective. We termed these 35 VRVs that were uniquely identified in Clusters TP2, TP3, and TP5 “pressing VRVs”.

### Comparison with AA Positions where Substitutions Have Been Identified Within US-Circulating VOIs and VOCs

We next compared the 90 VRVs and the 35 pressing VRVs with the 12 and 13 AA positions that have been shown to harbor substitutions (AA-subs) within US-circulating VOIs and VOCs, respectively (*2*). The 90 VRVs included 9 and 8 AA-subs in VOIs and VOCs, respectively; the 35 pressing VRVs included 4 and 8 AA-subs in VOIs and VOCs, respectively (Fig. 1C), even though all VOI/VOC sequences were excluded from the current analysis. Notably, 25 of the VRVs that have not been previously identified as an AA-sub in a VOI or VOC appear to have emerging trajectories, demonstrating the potential of the SLS method to identify novel Spike AA positions that may warrant further investigation/observation.

Five VOI/VOC AA-subs (Y144, F888, V1176, H69, K417) were not identified as a VRV. Fig. S3 shows the state/territory-specific relative abundances over time for states/territories where substitutions were identified at these 5 positions (albeit without meeting the statistical significance criteria for identification as a VRV). Our data suggest that, individually, these AA positions may be of less interest in US.

### Timely Detection of Emerging VRVs

Timely detection of potentially fast-emerging VRVs, and conversely, identification of VRVs likely not of concern, are both important for informing public health guidelines and for influencing research priorities. Given the importance of timely detection, we use the first time when a Pmax of a VRV exceeds 10% as the first reportable time. For each out of the set of AA-subs within VOIs/VOCs that were also identified as VRVs, Table 1 compares within each state/territory the time of detecting an emerging VRV as calculated by the SLS method vs. the first appearance of the AA-sub in the scientific literature. The SLS identified emerging VRVs in an average of 207 days, vs 299 days (average of reported values in literature). E484, an AA-sub in the B.1.1.7, P.1, and B.1.351 variants, is an exception as it was not detectable in the US until day 370, when it was first detected as a VRV in Rhode Island.

**Table 1.**
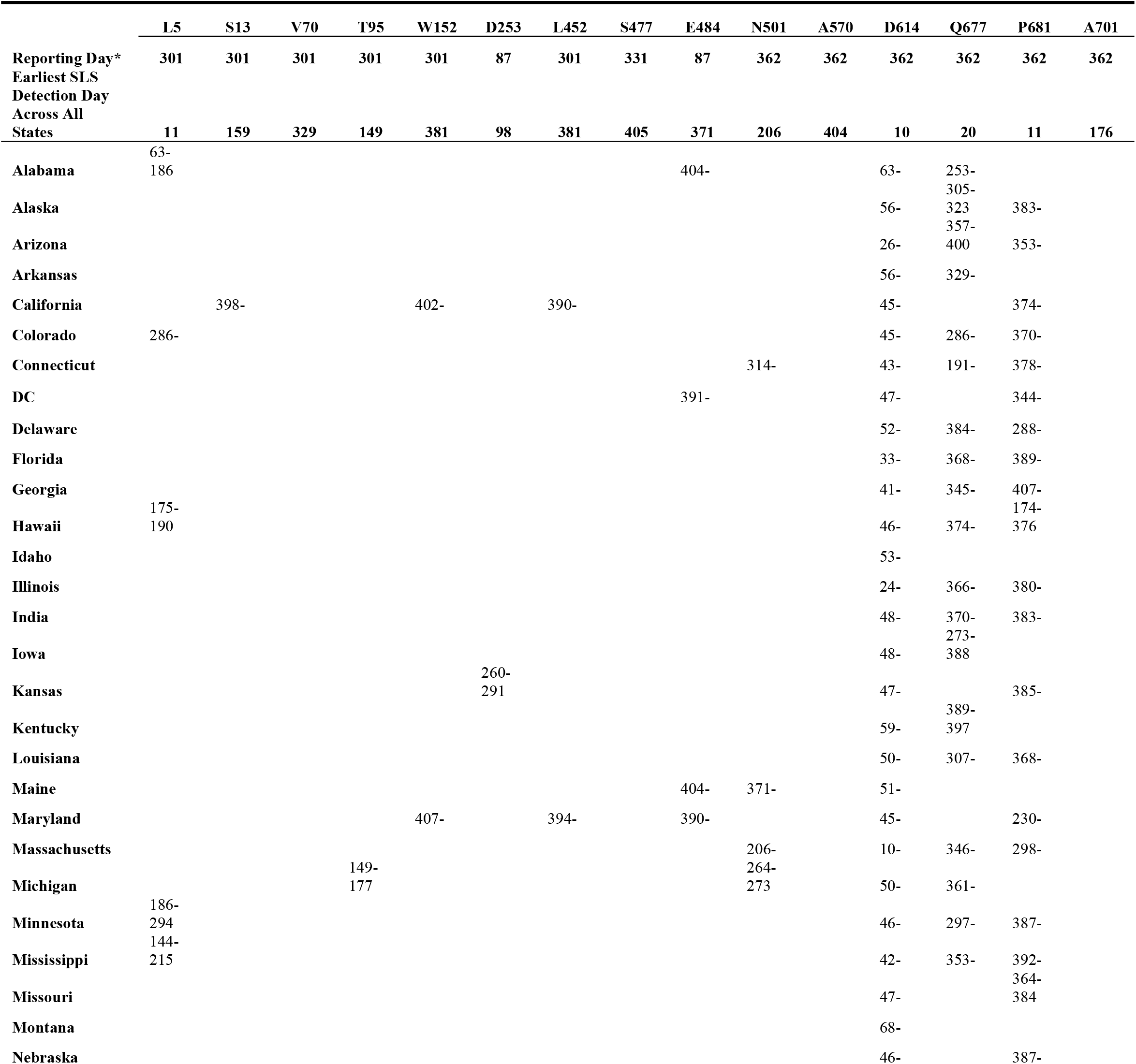
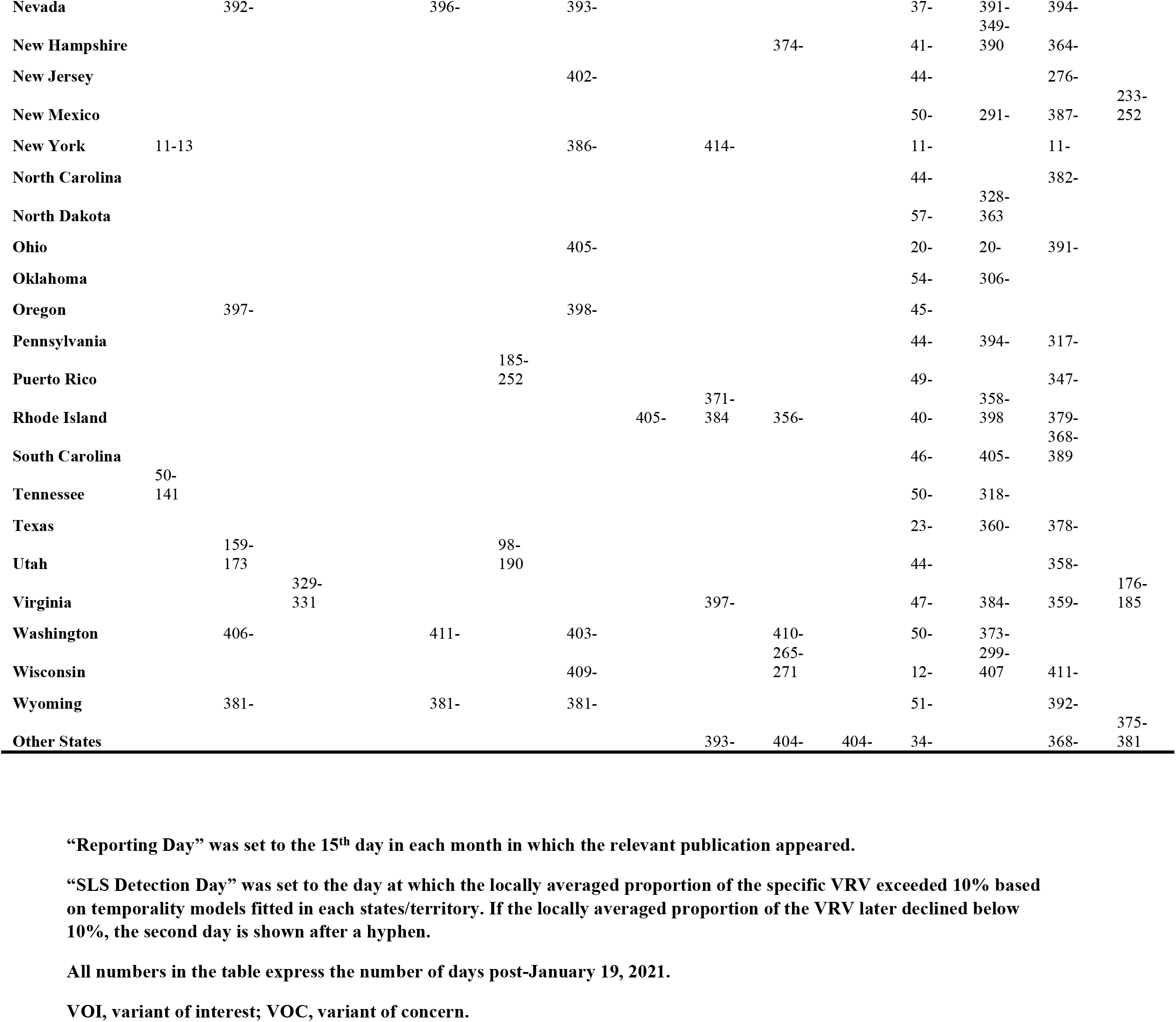
For 15 amino acid positions shown to harbor a substitution in a VOI or VOC, times estimated by the SLS method when the corresponding VRV had a locally averaged proportion exceeding 10% (and, if applicable, subsequently decreased below 10%) based on a state/territory-specific model. The top two rows show the first reported date in the literature of a VOI or VOC harboring a substitution at the designated site vs the date of VRV detection at the same amino acid position by the SLS method (across all states/territories).

### VRV-Haplotypes

SARS-CoV-2 is a single-stranded (“haploid”) RNA virus. The presence of multiple VRVs found in a patient form a VRV-haplotype. The accumulation of multiple VRVs on a single RNA strand could affect protein function more than a single VRV. To identify VRV-haplotypes, we performed unsupervised learning of selected VRVs and cases through a two-way hierarchical cluster analysis state/territory-by-state/territory. As shown in Fig. S4, some VRV-haplotypes are shared across states/territories, but most are not. Fig. 2, for example, shows the results of the unsupervised case and VRV clustering for Washington state. The heatmap shows that multiple VRVs tend to aggregate among subsets of cases, inspection of which can reveal VRV-haplotypes as follows: The case cluster “PG8”, which includes 14 cases, has VRVs from the “RG4” and “RG5” clusters, which include the VRVs (S13-W152-L452-V483-N501-D614-A684) (See Table S5).

**Fig. 2.**
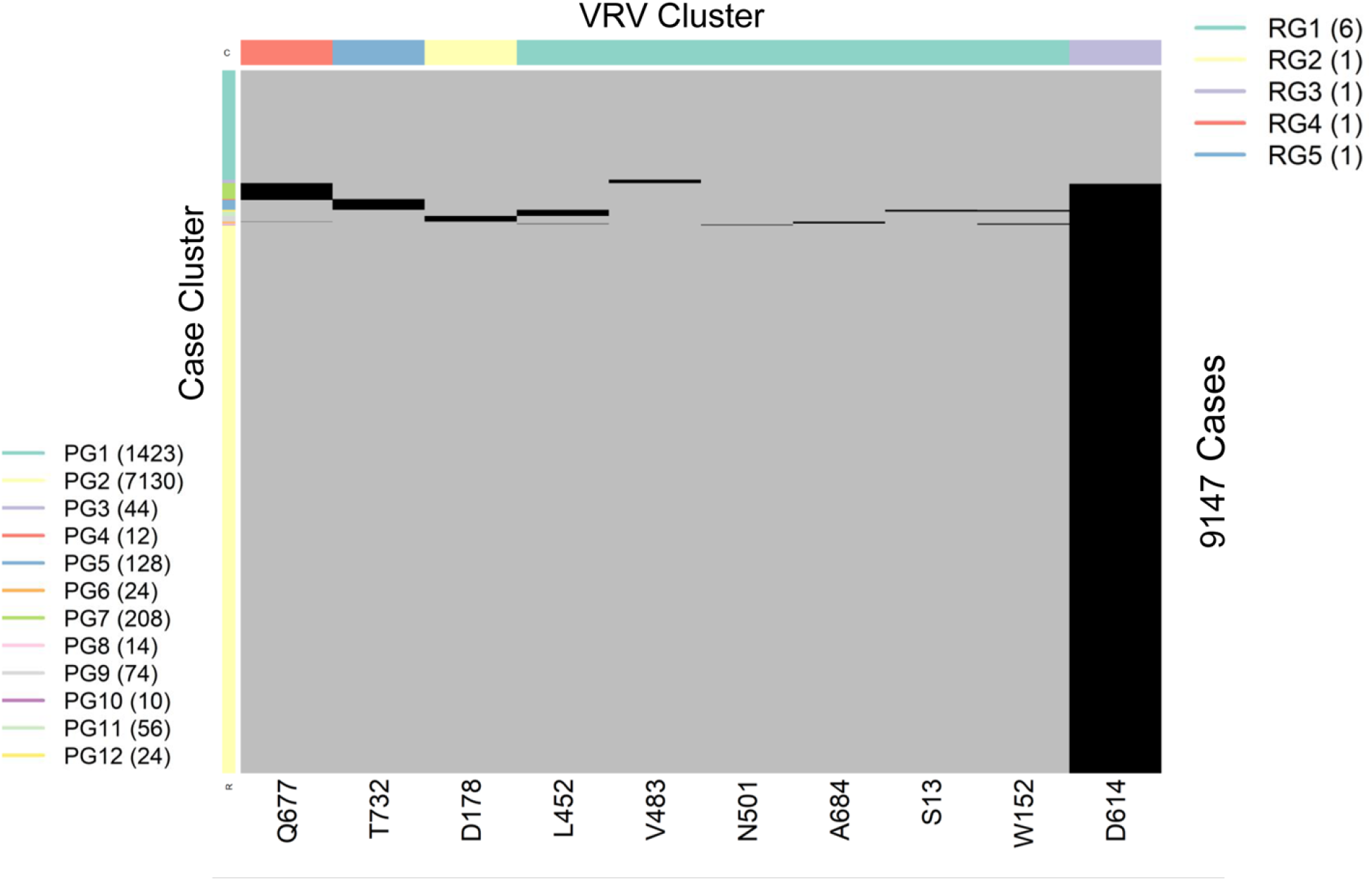
Heatmap showing the presence of 10 selected VRVs among 9147 cases in Washington state. Unsupervised learning was used to organize the 10 VRVs into 5 residue groups (RG1 through RG5) and to organize the 9877 cases into 12 patient groups (PG1 through PG12).

Collectively, these four case clusters were combined to identify the VRV-haplotype W1 (Table 2), found in 104 cases in Washington. Similarly, the case cluster “PG4” (12 cases) had three VRVs (D614, Q677, T732) from the “RG3”, “RG4”, and “RG5” clusters. In total, six VRV-haplotypes (W1 through W6) were identified in Washington, while the “W6” cluster (7130 cases) carried only a single VRV, D614 (Table 2). Comparison across VRV-haplotypes suggested that W6 evolved to W3, W4, and W5 via the acquisition of an additional mutation at T732, Q677, and D178, respectively. Similarly, both W3 and W4 could have evolved to W2 via the acquisition of an additional mutation at Q677 or T732, respectively.

**Table 2.**
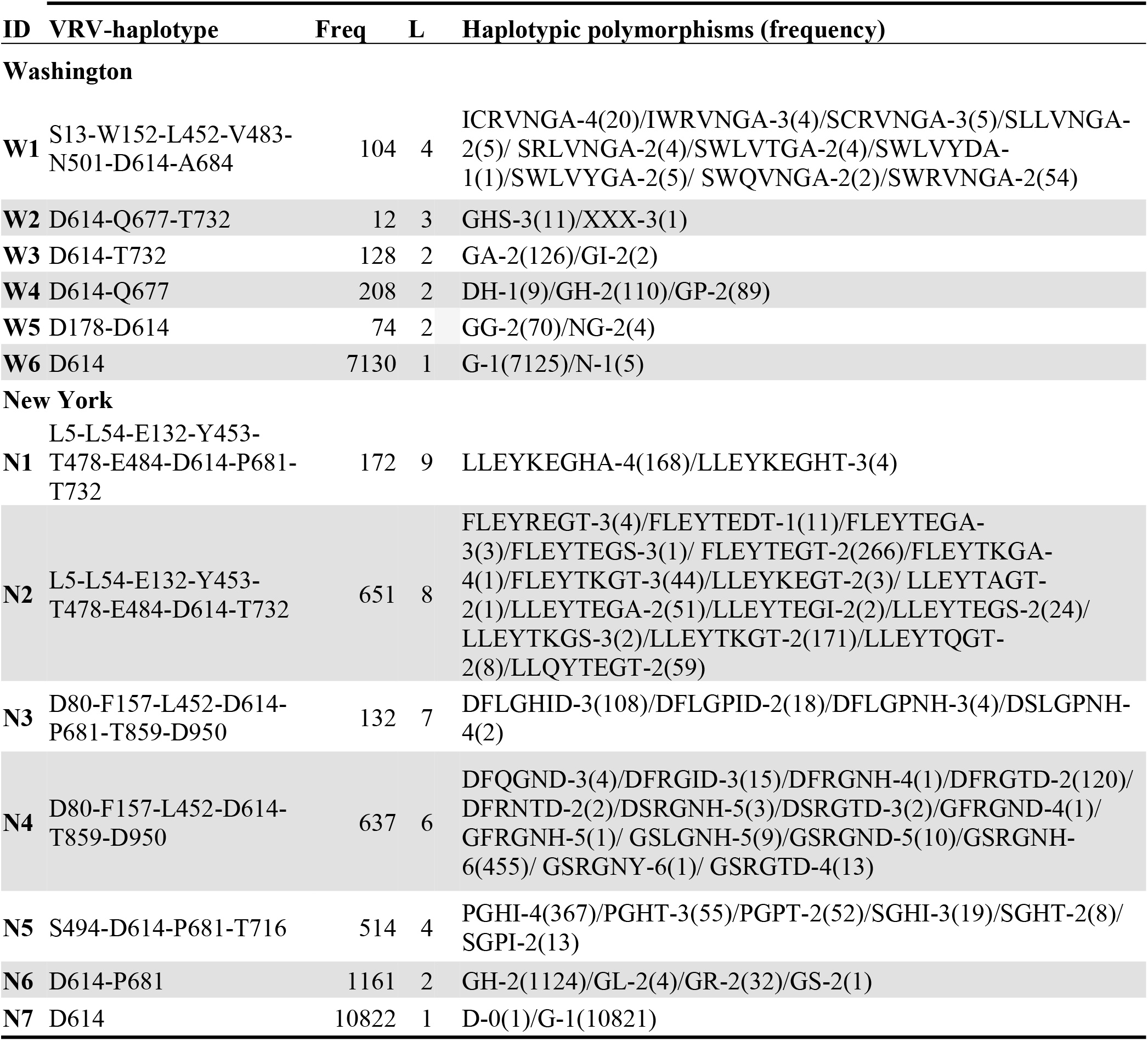
VRV-haplotypes identified in Washington and in New York: state-specific frequencies of cases, number of VRVs per VRV-haplotype, and haplotypic polymorphisms (state-specific frequencies). Unimputable residues are denoted with an “X”.

VRV-haplotype blocks are identified from unsupervised learning. Within each block, there can be multiple VRV-haplotypes that consist of polymorphic residues; individual VRVs may take either the reference residue or a substitution. For example, VRV-haplotype W1 had 10 haplotypes (Table 2), where the number after the hyphen indicates the number of substitutions. For example, the haplotype “ICRVNGA” has four substitutions, and was observed twenty times in Washington.

Table 2 also displays the VRV-haplotypes observed in New York (N1 through N7). The most frequent block, N2, has seven VRVs and 16 unique haplotypes. Block N1 only differs from Block N2 via the acquisition of the P681 VRV, and thus the two blocks are closely connected. Similarly, Block N4, which probably gave rise to Block N3, has 14 unique haplotypes, including “GSRGNH” (six substitutions), which was observed 455 times. Lastly, N5 probably arose from N6 via N7, and has the “PGHI” haplotype (observed 367 times). We next used unsupervised learning to construct haplotypes in Washington and New York of the 35 pressing VRVs (Table S6).

### Naming VRV Haplotypes via PANGO Lineages

As all sequences corresponding to VOI/VOC were excluded, the strains with detected VRVs are not currently undergoing special monitoring or characterization. We were thus interested in naming identified VRV-haplotypes and the PANGO lineages assigned by GISAID. To this end, we selected VRV-haplotype blocks including 4 or more pressing VRV mutations, resulting in 8 VRV-haplotype blocks. Table 3 cross-tabulates these VRV-haplotypes by their assigned lineages. Of particular interest, viruses with the haplotype “KGHA” of T478-D614-P681-T732 were observed 2132 times, and 2029 of them were assigned to the strain B.1.1.222. It is natural to name the haplotype T478K-D614G-P681H-T732A as a B.1.1.222. Another noteworthy strain is B.1.234, which corresponds to “SVGHF” and “SVGHS” of G142-E180-D614-Q677-S940 with exceptionally high frequencies (353 and 262). The remaining VRV-haplotypes mostly correspond to B.1. Fourteen other strains were found in more than 10 occurrences and may also be of potential interest.

**Table 3.**
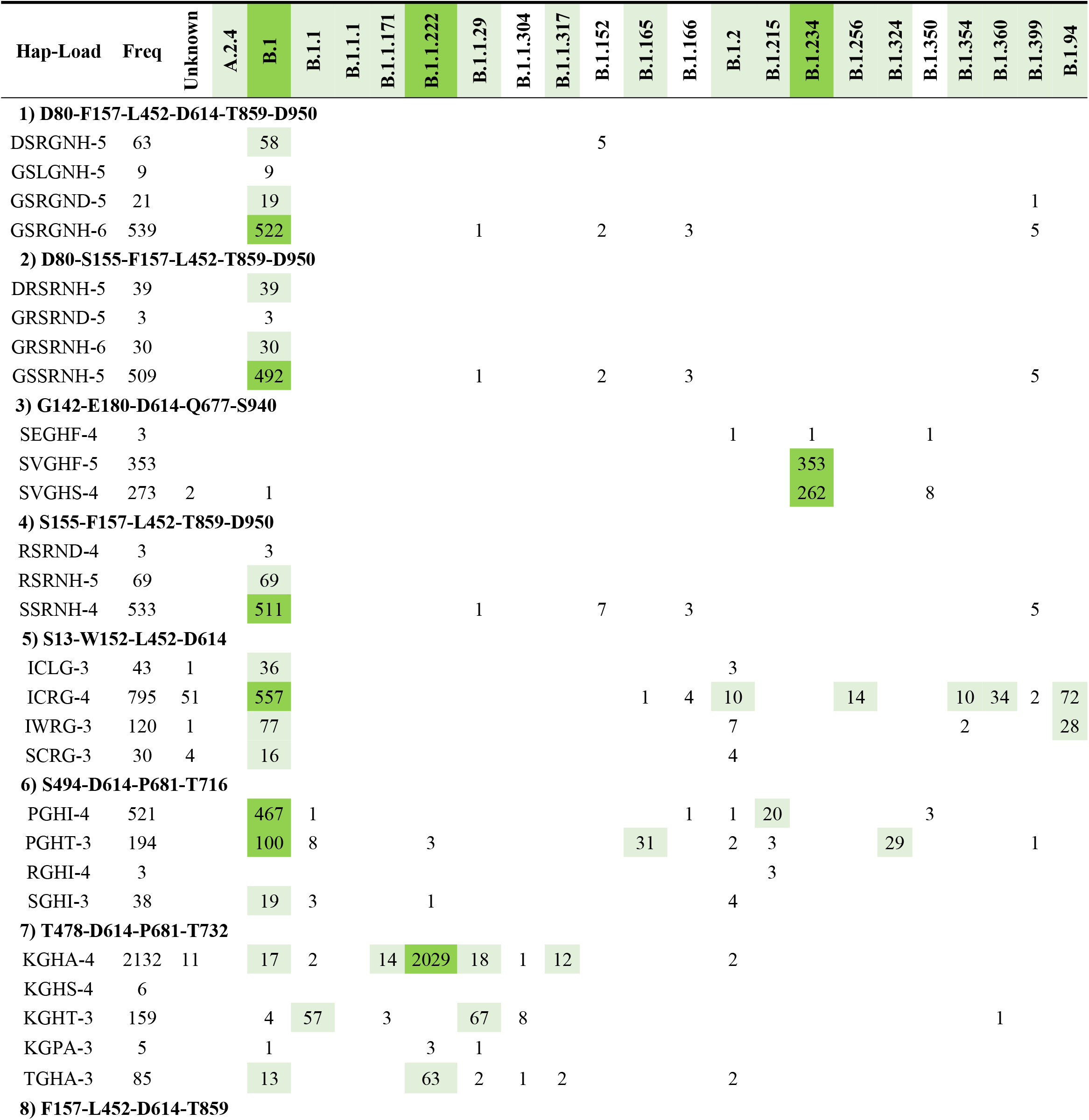
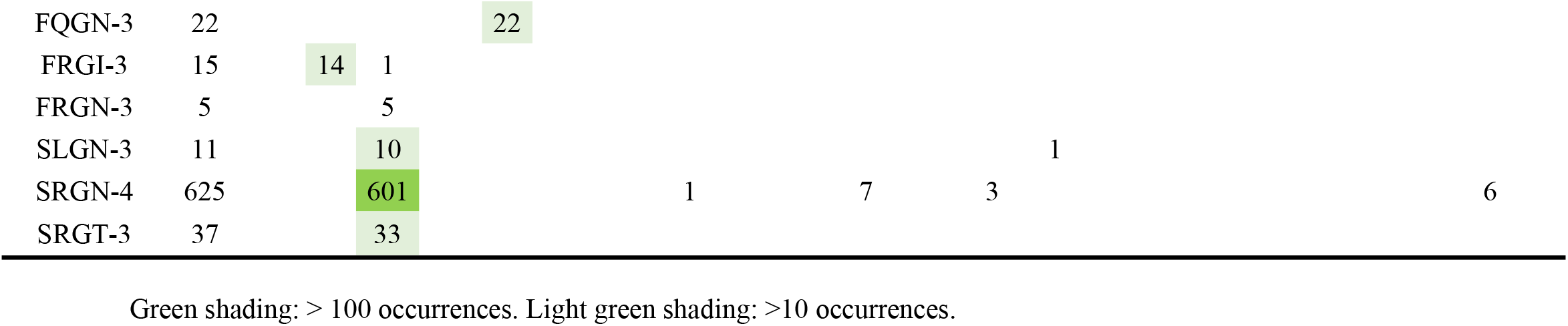
VRV-haplotypes. Cross-tabulation of individual VRV-haplotypes with GISAID-assigned lineages in all 167,893 sequences, excluding lineages with fewer than 10 occurrences. “Freq”, corresponding haplotype frequencies; “Unknown”, sequences not assigned to any image.

### Impact of VRV Haplotypes on Viral Structure

The SLS method includes homology modeling of Spike mutations, to predict possible consequences on Spike structure/function and to guide laboratory research. Inspection of the temporal dynamics of the VRV-haplotypes may be useful for identifying VRVs of interest. We performed homology modeling on two potentially interesting VRV-haplotypes, W1 (N501-A570-D614-P681-T716-S982-D1118, from the UK variant cluster B.1.1.7) and W2 [S13-W152-L452-D614, from the US variant cluster (B.1.94; B.1.427; B.1.429)].

The D614G mutation observed in the W1 haplotype has been associated with increased infectivity/transmissibility (*23-25*). Cryo-electron microscopy structures have been reported recently (*26, 27*) that reveal the structural consequences of this mutation and provide a plausible mechanistic explanation for the increased infectivity of D614G-carrying variants. The D614 VRV has predominated in all US cases for which sequence information is available in the TP2 cluster (Fig. 1B, Table S2). The N501Y mutation (present in the B.1.1.7 variant) is located in the receptor-binding domain (RBD) and has been reported to enhance binding affinity to the angiotensin-converting enzyme-2 (*10, 28*). N501Y has also been shown to reduce susceptibility to some nAbs, although the B.1.1.7 variant appears to remain susceptible to some extent to natural infection-acquired and vaccine-induced nAbs (*10*).

Of the five remaining VRVs in the W1 haplotype, A570, T716, and S982 seem relatively benign in that mutations at these positions are already decreasing in certain states/territories (this trend is also true to some extent for N501Y). While this observation may simply reflect inadequate sequencing efforts in recent months, it may also indicate that mutations at these positions do not confer any fitness advantage to the virus.

The two remaining VRVs in the W1 haplotype, P681 and D1118, are more intriguing. Mutations at these two sites, particularly at P681, appear to persist in multiple states/territories. The P681H mutation occurs in the S1/S2 cleavage segment of the Spike protein, which is typically not resolved in cryo-electron microscopy or x-ray diffraction experiments. Thus, we cannot speculate on potential structural consequences of this mutation. However, the continued presence of this mutation in many states and its location in the Spike protein S1/S2 cleavage segment suggest that it may warrant further investigation. We are not aware of any reports that D1118H impacts transmissibility or morbidity, but the location of this mutation in the Spike protein trimer assembly (Fig. 3A, B) suggests it could impact trimer assembly structure/stability/dynamics.

**Fig. 3.**
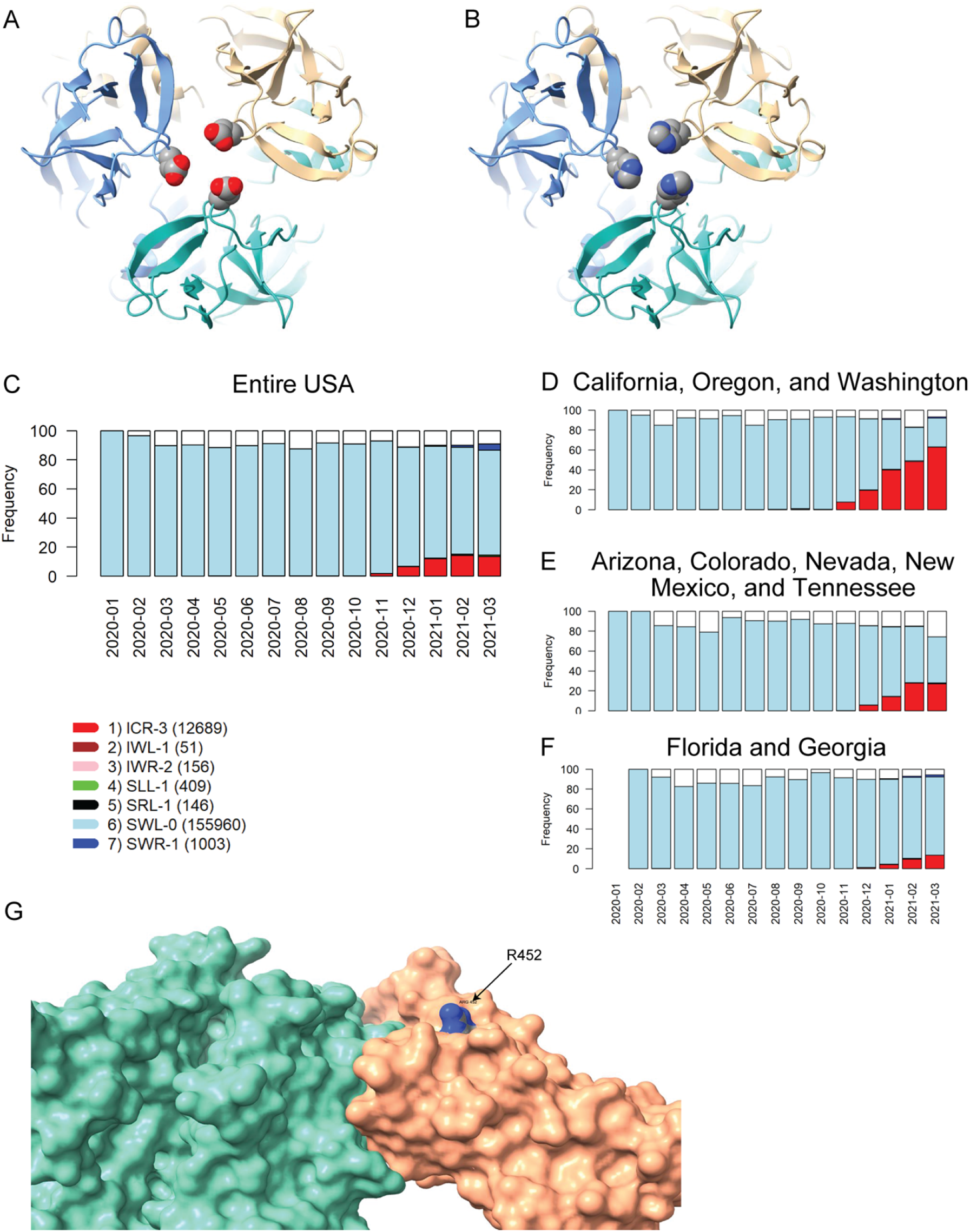
Homology modeling of Spike mutations and haplotypic polymorphisms over time of the S13-W152-L452 VRV-haplotype. (**A, B**) Modeled structure of the Spike protein trimer with (A) D1118 or (B) H1118 (homology-modelled using PDB entry 7KRS as the template structure). Spike protein monomers are displayed in blue, salmon, and aquamarine; aspartic acid and histidine residues are rendered as CPK images. (**C – F**) Frequencies over time for seven commonly observed haplotypic polymorphisms of the S13-W152-L452 VRV-haplotype, out of its polymorphisms in the US. Only haplotypic polymorphisms with at least 50 observations are included. Nomenclature is as follows: The first three letters designate the amino acids present at positions 13, 152, and 452, respectively; the number after the hyphen designates the number of amino acids at these three positions that do not match their reference strain equivalents. Numbers of sequences harboring each S13-W152-L452 haplotypic polymorphism (across the entire USA) are shown in parentheses. Frequencies of seven common S13-W152-L452 VRV-haplotypic polymorphisms (**C**) in the entire US; (**D**) in California, Oregon, and Washington combined; (**E)** in Arizona, Colorado, Nevada, New Mexico, and Tennessee combined; and (**F**) in Florida and Georgia combined. (**G**) Homology-modeled complex of the receptor-binding domain of the Spike protein (salmon), harboring the L452R mutation, bound to the angiotensin-converting enzyme 2 (ACE2) receptor (aquamarine). Within the R452 residue, nitrogen atoms are shown in blue and carbon atoms are shown in grey.

The US variants also carry the D614G mutation. The VRV-haplotype S13I-W152C-L452R (ICR-3) appeared in Fall 2020 and is rapidly becoming dominant in states on the West Coast, as well as appearing in selected Southwestern and Southeastern states (Fig. 3C-3F). The S13I and W152C mutations, which are situated in the N-terminal domain (NTD) of the Spike protein, have been implicated in escape from NTD-targeting monoclonal antibodies (*29*). The L452R mutation is situated in the RBD; homology modelling of the RBD-ACE2 complex shows that while R452 does not directly contact ACE2, the guanidinium side chain of R452 is surface-exposed and thus could potentially impact nAb binding (Fig. 3G). The L452R mutation was recently shown to reduce binding affinity to some RBD-targeting monoclonal antibodies, as well as to reduce susceptibility to nAbs (*29*). Thus, structural modeling of mutations in the S13-W152-L452 VRV-haplotype yields results consistent with the temporal dynamics of this VRV-haplotype.

## DISCUSSION

The continuous evolution of SARS-CoV-2 has already impacted public health guidelines and research priorities, with the potential of even more clinically consequential variants still to emerge. Here we leveraged a public data resource and described a statistical learning strategy for analyzing large, complex SARS-CoV-2 sequence datasets while incorporating temporal and spatial information. We provide detailed information on the emergence and persistence (or disappearance) of specific mutations in US states/territories, helping identify mutations that may warrant further observation/investigation. Our approach can be applied to other pathogens for which sufficient genomic surveillance data are available, generating important, statistically rigorous, and visually interpretable information for the biomedical research community, clinicians and public health officials. Our approach can also provide insight on the evolution of mutants and linkage with known viral strains.

By applying the SLS method to 167,893 US sequences not classified as any VOI/VOC, we identified 77 novel individual VRVs, including 25 pressing VRVs that appear to have emerged in the US. Among these pressing VRVs, the haplotype (T478-D614-P681-T732) links with the strain B.1.1.222 and (G142-E180-D614-Q677-S940) with the strain B.1.234, both of which do not correspond to any current VOI/VOC. Also of note, if the SLS method is applied to all US sequences, all circulating VOI/VOC are identified (results not shown).

As part of the assessment of immune correlates of protection, many randomized, placebo-controlled COVID-19 vaccine efficacy trials measure Spike protein sequences from symptomatic COVID-19 endpoint cases, and sometimes also from SARS-CoV-2 asymptomatic infections. Sieve analysis of these viral sequences can be conducted to assess whether and how vaccine efficacy depends on Spike protein sequence features, including differential vaccine efficacy across the levels of VRVs and of VRV-haplotypes (*30*). The graphical tools proposed here for spatiotemporal tracking of VRVs and VRV-haplotypes can be useful for sieve analysis, first by helping define and communicate the set of VRVs and VRV-haplotypes of study endpoint cases that have sufficient variability to be able to assess whether vaccine efficacy depends on the feature. For example, given that most vaccines use the Wuhan strain as the vaccine-insert, VRVs that meet our Pmax > 0.10 criterion would readily have the level of variability required for sieve analysis, whereas VRVs with Pmax < 0.02 would likely not. Secondly, including assignment to vaccine or placebo as a factor in the unsupervised clustering graphics applied to the vaccine efficacy trial sequence data sets may help communicate results of sieve analysis. Third, many of the vaccine efficacy trials have been offering the vaccine to placebo recipients, such that the placebo arm is lost and long term follow-up occurs only in individuals originally vaccinated or newly (deferred) vaccinated (*31*). The graphical tools may be applied to track study participant vaccine breakthrough virus VRVs and VRV haplotypes over time, and to similarly track VRVs and VRV haplotypes in GISAID data bases of unvaccinated persons matched by geography and time, and a comparison of these two tracking results may aid sieve analysis during the long term follow-up period of the vaccine efficacy trials.

Evidence is mounting that neutralizing antibodies acquired by natural infection (*32, 33*) or through vaccination (*34, 35*) are a correlate of protection against COVID-19. Therefore, it will be critical to assess whether and how VRVs and/or VRV-haplotypes in the infecting strains impact neutralizing antibody titers attained by natural infection (*36*), as well as whether and how they impact neutralization sensitivity to vaccine-induced neutralizing antibodies (*12*) and/or monoclonal antibodies (*37*). One possibility is that the graphical tools used here could annotate VRVs and VRV-haplotypes according to impact on neutralization. Moreover, a subset of sieve analyses is designed to restrict to VRVs and VRV-haplotypes that are known to impact neutralization response to the given vaccine under study, to improve power and to contribute to understanding neutralizing antibody-based correlates of protection. Applications of pinpointing VRVs or VRV-haplotypes that impact vaccine efficacy, and to quantify their impact, include informing models for predicting vaccine efficacy against circulating virus populations, and to aid optimization of vaccine strain selection.

A limitation of our approach is that it is constrained by intrinsic sampling limitations, since all sequences were collected and contributed by laboratories without consistent sampling protocols. Hence, despite the large size of our dataset, the analyzed sequences were not nationally representative. Further, it is important to interpret our results in terms of VRV proportions among reported sequence data, rather than incidences or prevalence of VRVs, in the absence of reliably estimated denominators. To overcome this limitation, public health agencies need to consider a uniformly developed surveillance protocol, to sequence COVID-19 cases from well-defined populations.

## MATERIALS AND METHODS

### Spike AA Sequences

Spike AA sequences (genome position: 21563-25384) from 189,727 COVID-19 cases in the US and selected US territories, along with their associated metadata, were retrieved from GISAID (*38*) (https://www.gisaid.org/) on March 23, 2021. Geographic origin (one of the 50 US states, Washington DC, Puerto Rico, or the Virgin Islands) was available for 189,284 of the sequences. For 443 of the cases, no US state/territory origin information was available. To ensure adequate sample size, Spike sequences from North Dakota, South Dakota, and the Virgin Islands were combined with these 443 sequences, forming an “Other States” category (728 sequences). Among them, 21,391 sequences were classified as a VOI or VOC (Table S1). These sequences were excluded, leaving 167,893 sequences for the analysis (see Table S2 for monthly case numbers by state/territory).

### Sequence Alignment and Transformation to VRV Indicators

Spike protein sequences were aligned to the Wuhan reference sequence (*39*) using MAFFT (*40*), yielding a complete “rectangular residue sequence matrix”. Sequences with at least one AA mutation (compared to the reference) were identified, enabling transformation of the residue sequence matrix to a matrix of binary VRV (mutant) indicators. Monomorphic residues led to columns of zeros and were eliminated from further analysis. We use VRV in this work to refer to a single AA position that harbors a substitution. We reserve the term “variant” in this work for identified VOIs and VOCs.

### Statistical Learning Strategy (SLS)

#### Modeling VRV Temporal Dynamics

To model non-linear temporal dynamics, a generalized additive model (GAM) was used to regress the VRV indicator over sample collection time through a non-parametric regression model. Further details are given in the Supplementary Materials.

#### Visual Representation of Temporal Dynamics

Within-state/territory: Temporal dynamics of <8 VRVs within a given state/territory were visualized with a line plot. For visualizing temporal dynamics of ≥ 8 VRVs within a given state/territory, unsupervised learning was applied, grouping VRVs with similar temporal patterns. Results were visualized with a heatmap.

Spatially integrated: To visualize spatiotemporal VRV dynamics, all state-specific temporal dynamics were integrated and unsupervised learning (one-way hierarchical clustering with the Euclidean distance with weights in favor of recent temporal trajectories and the “ward.D2” agglomeration method) (*41*) was applied.

#### Missing Residue Imputation

Imputation of missing amino acid information is described in the Supplementary Material.

#### VRV-Haplotypes

A viral strain harboring multiple VRVs is referred to as a “VRV-haplotype”. To identify VRV-haplotypes, unsupervised learning was used to organize both cases and VRVs through a two-way hierarchical analysis (*41*). Further information is given in the Supplementary Material.

#### Homology Modeling of Selected Haplotype Mutants

After identifying specific Spike protein mutants of interest from VRVs and related VRV-haplotypes, standard homology modeling methods were applied to generate 3D models. Further information is given in the Supplementary Material.

## Supplementary Material

### Materials and Methods

Fig. S1. For all Spike residues with sufficient variation, scatterplots of the maximum proportion (Pmax) of sequences from a given state/territory harboring a mutation at a given amino acid position vs. q-value. Points in red represent residues that meet both criteria for classification as a VRV. Points in black represent residues that do not.

Fig. S2. Temporal patterns of VRVs identified in each state/territory.

Fig. S3. Locally averaged proportions over time for substitutions at 3 AA-subs in a VOI (Y144, F888, V1176) and at 3 AA-subs in a VOC (H69, Y144, K417) that were not detected by the SLS method in states/territories where at least three sequences had a substitution at the designated AA position. AA-sub, amino acid that has been shown to harbor a substitution in a US-circulating VOI or VOC. VOI, variant of interest; VOC, variant of concern.

Fig. S4. Presence of VRVs among all cases in each state/territory. A gray cell means the VRV was not identified in the given case; a black cell means that it was. Both cases and VRVs were clustered by two-way hierarchical cluster analysis.

Table S1. Distribution of the 21,391 VOI/VOC sequences by specific variant and by state/territory.

Table S2. Distribution of the 167,893 SARS-CoV-2 sequences by state/territory and by GISAID submission month, along with state/territory-specific distribution of the 21,391 VOI/VOC sequences that were excluded from the analysis.

Table S3. The 10 identified geo-VRV clusters (TP1 through TP10), based on temporal profiles. Table S4. Frequencies of the 90 viral residue variants (VRVs) by state/territory, from an unsupervised learning from bi-clustering of all States and VRVs.

Table S5. VRV-haplotypes identified within each state/territory, along with state/territory-specific frequencies. The “positivity” column indicates the proportion of mutations in each haplotype block.

Table S6. Identified haplotypes of pressing VRVs in Washington and New York: frequencies, numbers of VRVs and haplotypic polymorphisms (frequency) in each state.

## Funding

This study was supported by the National Institutes of Health/National Institute of Allergy and Infectious Diseases (https://www.niaid.nih.gov/) through award UM1 AI068635 to PBG.

The funders had no role in study design, data collection and analysis, decision to publish, or preparation of the manuscript.

## Author contributions

Conceptualization: LPZ, TL, PBG, TRH, JTS, LS, THP, DEG, KRJ

Methodology: LPZ

Investigation: LPZ, TRH, JTS, LS, THP, DEG, KRJ

Visualization: LPZ

Funding acquisition: LPZ, PBG

Formal analysis: LPZ, TL

Data curation: JTS

Supervision: LPZ, PBG

Writing – original draft: LPZ, TL, PBG, JTS, LNC

Writing – review & editing: LPZ, PBG, TRH, JTS, LS, THP, LNC, DEG, KRJ

## Competing interests

The authors declare that they have no competing interests.

## Data and materials availability

All sequence data analyzed here are publicly available at GSIAD (https://www.gisaid.org/).

## Notes

### Competing Interest Statement

The authors have declared no competing interest.

